# Early-exposure to new sex pheromone blend alters mate preference in female butterflies and in their offspring

**DOI:** 10.1101/214635

**Authors:** Emilie Dion, Li Xian Pui, Antónia Monteiro

**Author notes:** Correspondence to ED and AM.

## Abstract

Insects use species-specific sex pheromone blends to attract members of the opposite sex which express the corresponding molecular receptors. Given this lock and key mechanism used for species identification and mate choice, it is currently not well understood how pheromone blends or receptor systems evolve. One possibility is that insects develop preferences for new sex pheromone blends via the process of learning, and that these learned preferences may be passed on to the next generation. We tested these hypotheses by exposing newly emerged *Bicyclus anynana* female butterflies to either wild type or to modified male sex pheromone blends. A few days later, we scored female mating outcome in a choice trial involving both male types. We also assessed the mating outcome of naïve offspring of females that underwent distinct odor learning trials to test for a potential inheritance of learned odor preferences. Naïve (parental) females mated preferentially with Wt-blend males, but females pre-exposed to new blends either shifted their preference to new-blend males, or mated equally with males of either blend type; the response depending on the new blend they were introduced to. Naïve daughters of females who were exposed to new-blend males behaved similarly to their experienced mothers. We demonstrate that females are able to learn preferences for novel pheromone blends in response to a short social experience, and pass that learned preference down to the next generation. This suggests that learning can be a key factor in the evolution of sex pheromone blend recognition and in chemosensory speciation.

**Significance statement:** While the diversity of sex pheromone communication systems across insects is well documented, the mechanisms that lead to such diversity are not well understood. Sex pheromones constitute a species-specific system of sexual communication that reinforces interspecific reproductive isolation. When odor blends evolve, the efficacy of male-female communication becomes compromised, unless preference for novel blends also evolves. We explore odor learning as a possible mechanism leading to changes in sex pheromone preferences. We show that preferences for new blends can develop following a short learning experience, and that these novel preferences can be transmitted to the next generation. To our knowledge, this is the first investigation of sex pheromone blend preference learning impacting mate choice and being inherited in an insect.

## Introduction

The evolution of sexual communication via pheromones is a fascinating area of evolutionary biology because changes in pheromones or their perception may lead to assortative mating, reproductive isolation, and eventually speciation. In most insects, sex pheromones are critical to the process of finding and selecting a mate (1, 2). The composition and relative proportion of the sex pheromone blend components are typically species-specific, and, together with the corresponding specific reception molecules, play a fundamental role in interspecific reproductive isolation (3, 4). Recent studies in Lepidoptera, for instance, support a key role of this chemosensory system in speciation, where both male and female pheromone preferences have diversified along with the evolution of the respective blends (5-8). However, there is still very little understanding of the mechanisms originating divergence in mate preferences for new pheromone blends.

Learning to prefer a novel mate signal early in life could be a possible mechanism driving the evolution of new pheromone communication systems. Learned preferences for novel mate visual signals were previously shown in several arthropod species. In particular, early exposure to new ornamentations in spiders (9), fruit flies (10), or butterflies (11) all led to shifts in mate preferences in sexually mature older individuals. These premating experiences have thus been proposed to play a significant role in reproductive isolation (12, 13). Similar to visual signal learning, odor learning is known to happen routinely in an insect’s life. For instance, honeybees learn pollen odors while foraging or after being exposed to pollen at an early age (14), and parasitoids learn the odors of their hosts when laying their eggs (15). Moths can also learn to associate a sex pheromone component with a food reward after a few proboscis extension conditioning trials (16). To date, however, there is no data on whether any insect can learn to prefer novel pheromone blends, via an early exposure, that results in a change in mating outcome. Learned pheromone preferences, over time, could eventually become genetically assimilated and fixed in a population, giving rise to populations of insects with novel pheromone blends and with specific sensitivity for those blends encoded at the genetic level.

A mechanism that could accelerate the process of genetic assimilation could be the transgenerational inheritance of acquired traits (17). Behavioral variations following an environmental experience have been hypothesized to be caused by epigenetic modifications that affect the expression of relevant genes, and which can be inherited through the germline (18). In particular, inheritance of learning and memory processes has already been shown in several species. Attraction of the nematode *Caenorhabditis elegans* to olfactory signals after exposure to these cues was shown to be passed-down to their naïve offspring for several generations (19, 20). A more detailed molecular mechanism of learned odor avoidance was discovered in mice, where deterrence towards an odor was shown to be transmitted to the next generation via the inheritance of a hypomethylated form of the odor receptor gene expressed in the olfactory system (21). These examples illustrate how learning to avoid or prefer an odor might be transmitted by epigenetic marks to the offspring via the germ line. If epigenetic modifications, such as silencing marks, alter patterns of gene expression for a few generations, shielding these regions from natural selection, then genetic mutations are free to also accumulate in these same regions, eventually stabilizing the phenotype that was originally only environmentally induced (22).

In order to contribute to this field of research we tested whether female butterflies can shift their mate preferences after being exposed to males with novel sex pheromone blends, and whether learned preferences can be transmitted to the next generation. We performed these experiments on *Bicyclus anynana* butterflies, the only Lepidopteran in which mate preference learning has been shown to take place (11). In particular, females can learn preferences for novel male wing patterns in males if they are exposed to them for a short period after emergence (11). This learned preference, however, only happens if these males express the correct pheromone blend (23). Three male sex pheromones (MSP) have been identified in this species: (Z)-9-tetradecenol (MSP1), hexadecanal (MSP2) and R6, R10, R14-trimethylpentadecan-2-ol (MSP3). They are produced after emergence in specialized wing glands (24, 25). In *B. anynana* of the wet season form, females are the choosy sex (26, 27). Young virgin females frequently reject courting mates before reaching sexual maturity, being exposed to male sex pheromones during this process (24, 28). This particular life history, thus, lends the performance of pheromone learning experiments in this butterfly ecologically relevant.

To decide on the type of blend manipulations to do for this experiment we investigated how pheromone blends vary across closely related species. In insects, new sex pheromone blends may evolve in their composition by loss or gain of single components, or by variation in the ratios of components (29). A comparative MSP blend study across *Bicyclus* species showed that close relatives vary both quantitatively and qualitatively in their MSP blend. Sympatric pairs of species display larger differences in component amount and identity than allopatric pairs, suggesting that the MSP blend is involved in pre-mating reproductive isolation in this genus of butterfly (5). We decided, thus, to vary the amounts of blend components in our odor learning experiment in *B. anynana*.

Our experimental design and questions were as follows: We created two New Blends (NB) by either preventing the release of MSP2 and reducing the amounts of MSP1 and MSP3, thus creating a “reduced” blend (called NB1); and by increasing the amount of MSP2, producing an “enriched” blend (called NB2). We exposed immature females to males with these new blends and to males with the respective control manipulations (called Wt1 or Wt2), and observed the mating outcome of the same females in the presence of the two types of male a few days later (Fig. 1). We tested whether 1) new blend males are less attractive to naïve females; 2) females learn to prefer males with new blends and; 3) learned female preferences for one of the blends are transmitted to their offspring.

**Figure 1.**
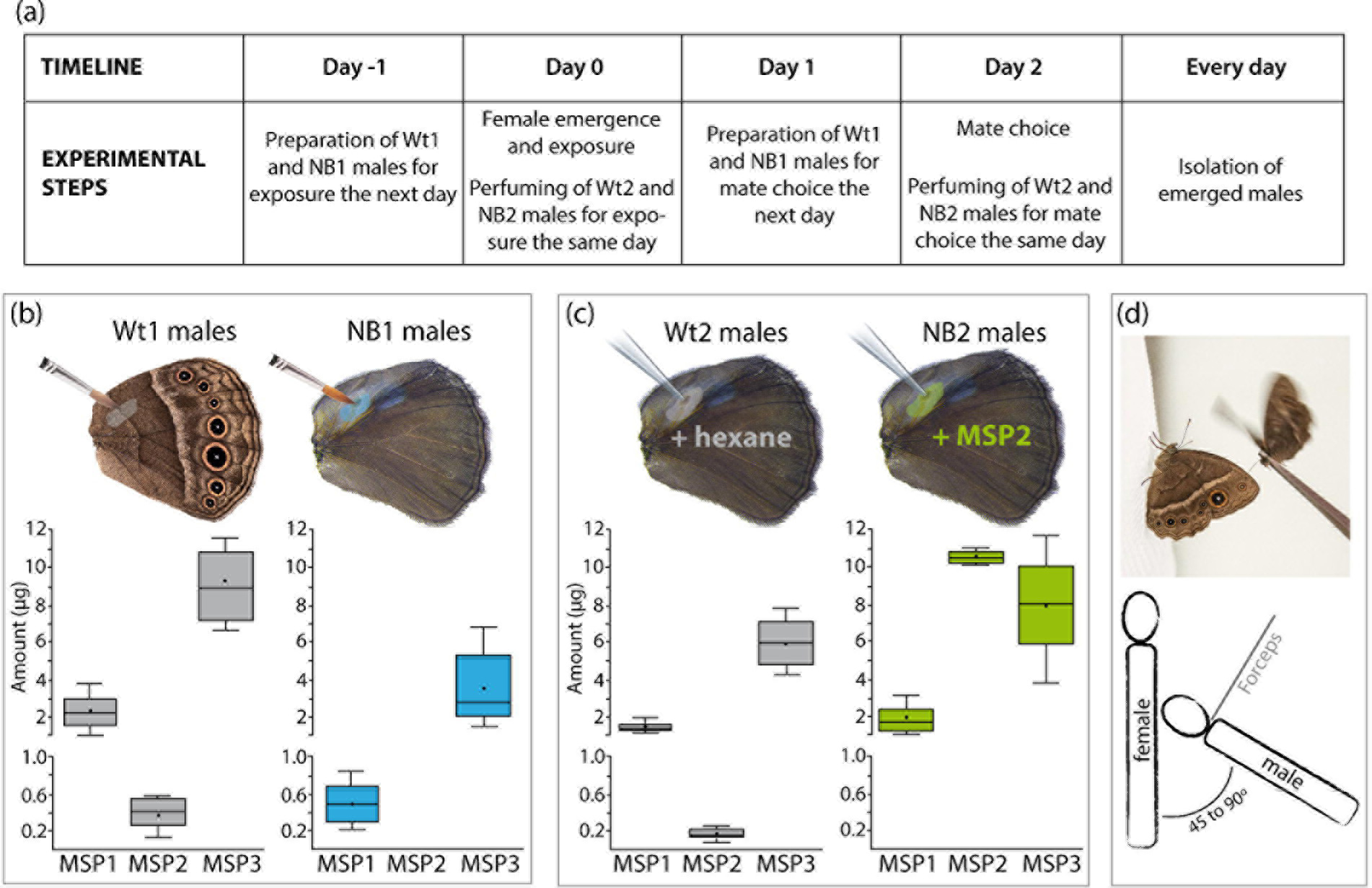
Experimental procedure. (a) The timeline of the experiment indicating when each step was performed. (b) Coating of the male androconia (NB1 males) prevented the release of MSP2 and reduced the total amount of MSP1 and MSP3 per male. (c) The average total amount of MSP2 per NB2 male, 30 minutes after perfuming with synthetic hexadecanal, is increased compared to Wt2 males. In each graph, the horizontal line and the point in each box are the median and the mean amount, respectively. The 25th and 75th percentiles are contained within the outline of the boxes, and the horizontal lines above and below each box show the 1.5 times inter-quartile range of the data. 5 to 10 males were used to measure MSP amount in each treatment. (d) Schematics of the female exposure where the bottom panel illustrates the position of both male and female individuals from a top view.

## Results

### Male manipulations altered the levels of MSPs

MSP2 was previously suggested to be the most relevant pheromone to female choice because males producing higher absolute or relative amounts of MSP2 had a higher mating success (30). By blocking the pheromone gland on the male hindwing or by perfuming the wing with MSP2, we created two novel blends, NB1 and NB2 respectively, that were different from Wt1 and Wt2 control blends (Fig. 1). In particular, MSP2 was absent in NB1 males, and was increased by 50 fold in NB2 males (30 minutes after perfuming) (Figs. 1b, 1c and Supplementary Fig. 1). Total amounts of MSP1 and MSP3 were reduced by an average of 70% and 60% respectively in NB1 males, compared to Wt1 males (Fig. 1b).

### New blend males are not attractive to naïve females

The innate sex pheromone preference of females without any social experience was monitored in a mate-choice assay, where the identity of the male (NB1 versus Wt1; or NB2 versus Wt2) that mated first with that female was scored. Naïve females showed an innate mating bias for the wild type blends in both experiments, with 77% and 70% of the tested females mating first with Wt1 and Wt2 males, respectively (Figs. 2a and 3).

**Figure 2.**
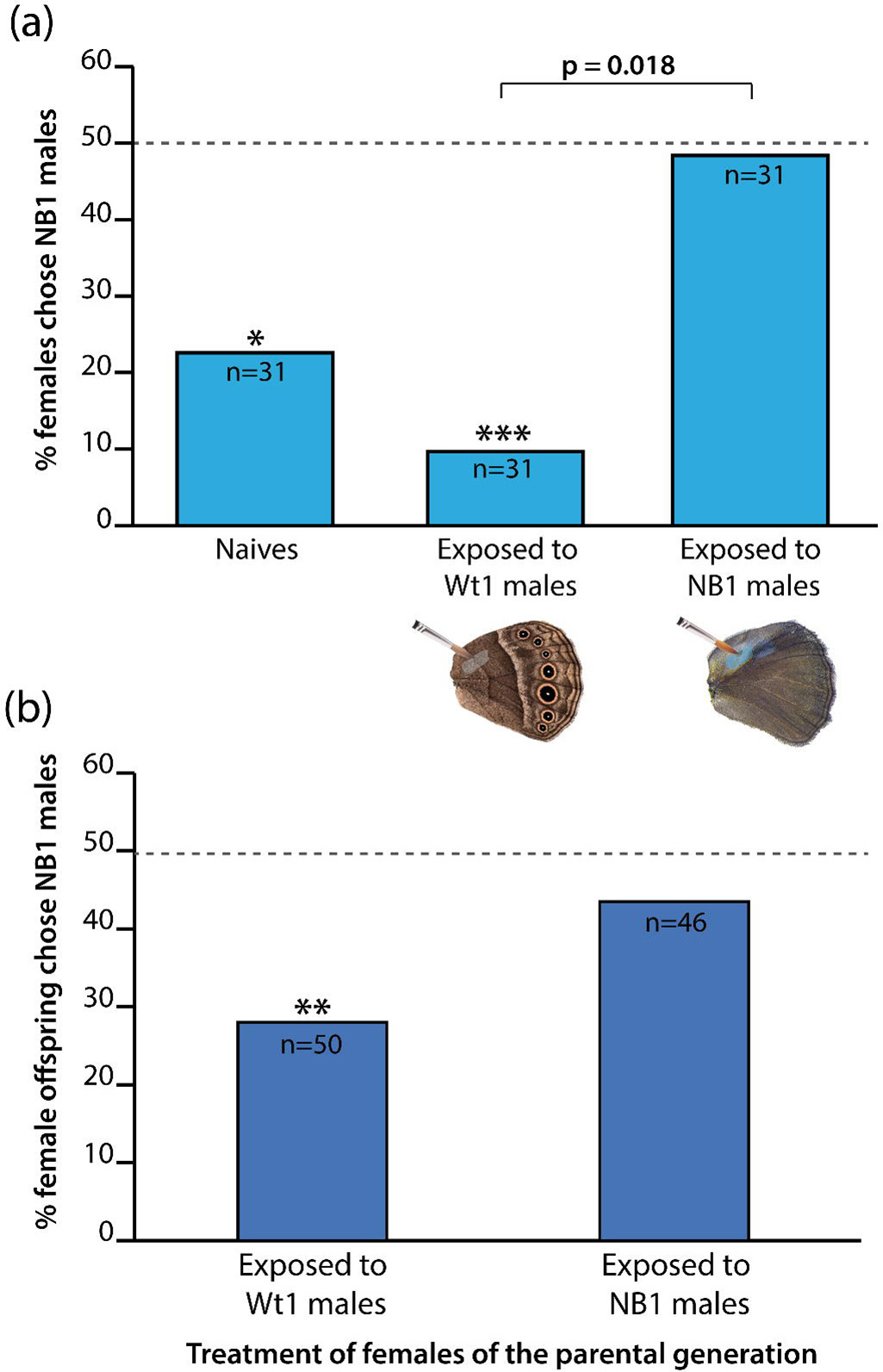
Mating outcome of females after exposure to the “reduced” (NB1) and wild type pheromone blends, and mating outcome of their naïve offspring. (a) Mating outcomes shifted after females were exposure to a male with a reduced blend. Most naïve females, and females exposed to Wt1 blend mated with Wt1 males, but females exposed to NB1 males mated with these males at significantly higher rates than Wt1-exposed females. (b) Offspring of females exposed to Wt1 mated preferentially with Wt1 males, similarly to naïve and Wt1-exposed females from the parental generation. However, offspring of NB1-exposed females mated equally with either male type. Asterisks (* p<0.05; ** p<0.01; ** p<0.001) indicate statistically significant preferences for the Wt1 blend using Pearson’s *χ*^2^ test. The dotted line at 50% illustrates random mating. The horizontal bar above the plot shows a significant difference in mating outcome between the two treatments (from the Tukey post-hoc test, adjusted p value is indicated). The “n” on each bar indicates the total number of female tested. Post-hoc test results providing adjusted p values comparing the different treatments are shown in Supplementary Table 1a.

**Figure 3.**
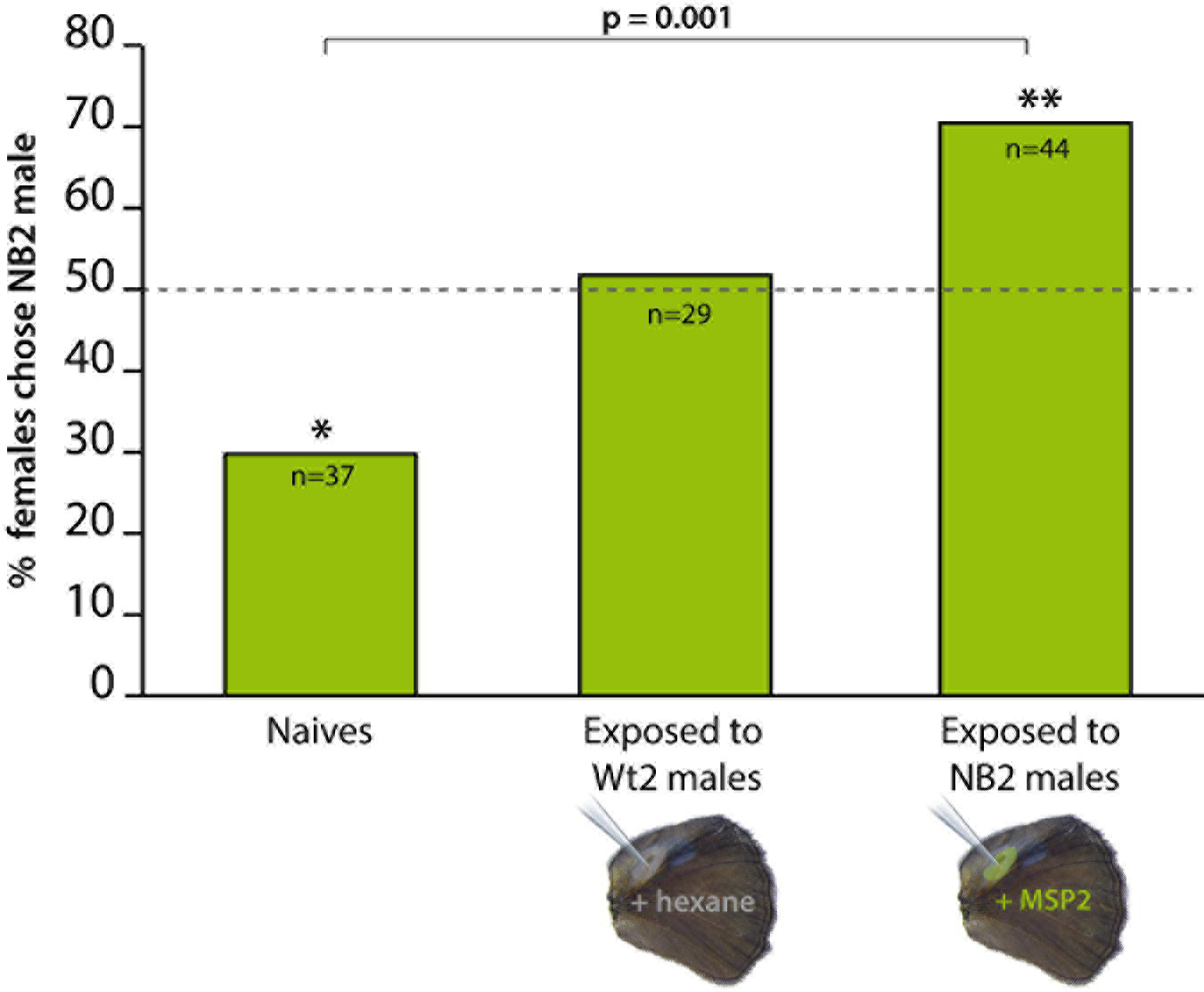
Mating outcome of females after exposure to the “enriched” (NB2) and wild type pheromone blends. In experiment 2, females shifted their mating outcome after exposure to a Wt2 male or a male perfumed with a novel pheromone blend containing more MSP2. Most naïve females mated with Wt2 males, females exposed to the Wt2 blend mated equally with both male types, and females exposed to the new blend mated with new blend males at significantly higher rates than naïve females. The dotted line at 50% illustrates random mating. Asterisks (* p<0.05; ** p<0.01) represent non-random mating outcomes using Pearson’s *χ*^2^ test. The horizontal bar above the plot shows a tested. Post-hoc test results providing adjusted p values comparing the different treatments are shown in Supplementary Table 1b.

### Premating exposure to novel MSP blends modified female innate mating bias

To test if female mating outcome changed after a short social experience, we exposed different females to either NB1, NB2, or to their corresponding control wild type males, and scored mating outcome in a mate choice assay a few days later. Because the males used for exposure and mate choice were from 4 to 6 days old, we used mixed models to measure the effects of these variable along with the effect of exposure treatment on female mating outcome (see method section for full details). Female premating exposure treatment significantly affected subsequent mating outcomes in both experiment 1 (MERL: *χ*^2^=16.7, Df=4, p=2.2 e^-4^) and experiment 2 (GLM, F= 6.7, Df=2, p=1.8 e^-3^). In particular, 90% of the females pre-exposed to Wt1-males mated with Wt1-males, showing a strong significant preference for the Wt1-blend, whereas females pre-exposed to NB1-males showed no mating bias, mating randomly with either male (only 51% mated with Wt1 males; Fig. 2a). These two mating outcomes were significantly different (Post-hoc tests from MERL, adjusted p = 0.018). In the “enriched blend” treatment (Fig. 3), females pre-exposed to Wt2-males showed no mating bias, 51% of them accepting the NB2-male first for mating. NB2-exposed females mated predominantly with NB2-males (70%), while naïve females mated predominantly with Wt2-males (70%). These two mating outcomes were also significantly different (Post-hoc tests from GLM, adjusted p=0.001).

### Female offspring had similar preferences as their exposed mothers

To test for inheritance of learned preferences, we submitted each naïve offspring of NB1-exposed and of Wt1-exposed females to mate choice trials with a single NB1 and a single Wt1 male. Note that the mothers of these female offspring, despite differences in early odor exposure, all mated with Wt males to control for this variable. Offspring of females exposed to Wt1 males mated preferentially with Wt1 males (72%; Pearson’s test: *χ*^2^=9.7, p=0.003), whereas offspring of females exposed to NB1 males did not show any mating bias, mating randomly with both male types (57% mated with Wt1-males; Pearson’s test: *χ*^2^=0.8, p=0.4), as did their mothers (Fig. 2b). The percentage of matings with NB1 males was 15% higher in offspring of NB1-exposed females than in offspring of Wt1-exposed females (Fig. 2b). For a difference of this magnitude (i.e., effect size) to be significant across offspring types, the sample size would need to be increased to an average of 275 tested female offspring in each group (Supplementary Table 2).

In all experiments, the age of males used for the pre-mating exposure, mating trial, and the position of the black dot placed on the wings to differentiate NB2 and Wt2 males (experiment 2) did not significantly affect mating outcome (Supplementary Table 3).

## Discussion

### Females learned to prefer a mutant pheromone blend

Naïve females mated preferentially with males with a Wt blend over males with either of the mutant blends tested. These results demonstrate the ability of the olfactory circuitry to distinguish the different blends and confirm that the specific male sex pheromone composition and ratios of components are important for *B. anynana* mate selection (24, 28, 30). We demonstrate, however, that an early and brief exposure of females to novel pheromone mutant blends alters their subsequent mating patterns. Initially unattractive males, lacking MSP2 and producing less MSP1 and MSP3, became as attractive as Wt males after a short early-exposure of females to their mutant blend. More strikingly, females mated preferentially with originally unattractive males with high amounts of MSP2, after they were exposed to this new blend. The changes in mating outcome are likely to have resulted from a change in female behavior rather than from alterations in male-male competition or male behavior during the mate choice trial due to the male’s different odors. This is because the mate-choice experimental set-up with both males was identical in every treatment. In addition, the shift in the butterflies’ mate preference was not influenced by mate-choice copying (31), as all females were isolated from each other and from the males since the pupal stage, and visually isolated from each other at every point in the experiment, including during mate choice trials. These results lead us to conclude that female preference for a male pheromone odor blend in *B. anynana* is not fixed but plastic, and influenced by early pheromone odor experiences.

Female preference learning was stronger towards NB2 than NB1 blends but it is still unclear why this was the case. In particular, NB2-exposed females preferred NB2 over Wt2 males, but NB1-exposed females only lost their preference bias towards Wt1 males, mating randomly with either male type. Previous work showed that males with either higher absolute or relative levels of MSP2 to other MSP components had higher mating success. MSP2 was, thus, proposed as the most relevant MSP to female choice (30). Here our data for naïve female mating outcome showed that females actually discriminate against males with very high levels of MSP2, but upon exposure to these high levels, females subsequently mate more frequently with these males. We propose that it might be harder for exposed females to overcome the unattractiveness of NB1 compared to NB2 because NB1 is a highly divergent blend lacking MSP2, whereas NB2 has increased amounts and relative ratios of MSP2. Another possibility for this asymmetry in learning, which will need additional testing in future, is that female exposure to enhanced blends (with additional components) relative to Wt blends, leads to overall stronger mate discrimination ability, whereas exposure to weaker blends relative to Wt, leads to loss of mate discrimination abilities. When females are exposed to low amounts or absence of components (as it is the case for NB1- and Wt2-exposed butterflies), they are less discriminatory and mate randomly with either male. However, when females are exposed to higher levels of blend components (such as Wt1-and NB2-exposed females), they discriminate between NB and Wt males, preferring the blend they have been exposed to. We also note that an increase in MSP2 amount alone (as in NB2 males) is sufficient to trigger a change in female discriminating abilities, confirming that this component is important in *B. anynana* mate choice. The neurological mechanisms involved in this process, however, are still unclear.

### Alterations of the chemosensory system may be responsible for the change in female blend preference

Brief exposures to odors were previously shown to impact the expression of olfactory receptors, odorant binding proteins, and the development of brain olfactory centers in honeybees and moths (32–35). In the bee, qRT-PCR analysis revealed that 6 floral scent receptors were differentially expressed in the antenna depending on the scent environment they experienced (32). A brief one-minute exposure of male moths to female sex pheromones led to the up-regulation of a pheromone-binding protein in the male antennae, an enlargement of the antennal lobe, and an increase in the volume of the mushroom bodies in the male brain, which resulted in a higher sensitivity of the exposed males to the female blend (33-35). The brief exposure of *B. anynana* females to the new male pheromone blend may have led to similar changes in the female brain. The mechanisms in place, however, require future exploration.

### Learning to prefer a mutant blend male may have important evolutionary consequences

Both empirical and theoretical studies have highlighted how the learning of a trait or a mate preference can impact assortative mating and population divergence (12, 36). Depending on the specific ecological conditions, type of trait, or learning process, models predict that mate preference learning can lead to reproductive isolation (*e.g*. (13, 37). Moth and butterfly sex pheromone blends are highly species-specific, ensuring the precise recognition of a compatible mate. These blends are generally thought to be under stabilizing selection because altered signals are less attractive and are thus selected against (29). However, the learning process that we describe here, by allowing males with divergent blends to reproduce, may mitigate the strength of stabilizing selection, and create opportunities for pheromone blends and reception systems to evolve. A recent study suggested that quantitative and qualitative variations observed in blends within and between natural *B. anynana* populations are potentially catalyzing ongoing speciation (6). The odor learning ability of *B. anynana* females has probably maintained the high variance in MSP amounts measured in different stock populations (24, 25, 30), as 225 well as the variance in MSPs detected across natural populations (6). Furthermore, the use of multimodal signals in mate selection in *B. anynana*, where females use both olfactory and visual signals to assess mate quality (11, 23, 28), may facilitate pheromone learning and the evolution of the MSP blends. The presence of species-specific visual cues on the male wings likely increases a female’s acceptance of odor-unattractive males from the same species, and decreases the risks of females learning new blends from hetero-specifics that could lead to hetero-specific mating. Thus, learning to prefer novel odors or odor blends may be a key starting point in the process of reproductive isolation and speciation, especially if this preference can be transmitted to the next generation via the germ line.

### Transgenerational inheritance of pheromone preferences may facilitate the evolution of assortative mating and speciation

Naïve female offspring of mothers exposed to NB1 blends stopped avoiding NB1 blends, as did their naïve mothers, indicating that habituation towards this new blend was transgenerationally inherited. Daughters of females exposed to Wt-blend males, however, did not increase their preference for Wt-blend males. This lack of transmission of a more extreme preference for Wt blends in female offspring could be explained by an exhaustion of genetic variation, since exposure of females to wild type butterflies has been repeatedly done presumably since the origin of this species. Because all F1 individuals were kept completely isolated from their conspecifics until mate choice, a change in F1 female preference is also not a result of social transmission, but more likely mediated via epigenetic mechanisms.

The transgenerational inheritance of acquired behaviors remains a controversial topic despite the growing number of empirical work supporting it, including mechanistic studies. For instance, first- and second-generation naïve Drosophila melanogaster offspring displayed a preference toward the alcoholic odors their parent where trained to like. Disruption of the F0 olfactory receptors and specific neuron inputs into the mushroom bodies abolished the change in offspring response, identifying potential targets of epigenetic transmission (38). In addition, a number of studies have revealed that DNA methylation regulates memory formation and learning processes in insects (e.g. in bees (39, 40)) but have not investigated whether these marks can be inherited to the next generation. Inheritance of a differentially methylated odor receptor gene, however, was shown to take place in mice that learned to avoid a specific odor (21). We speculate that in our system, genes involved in odor perception and/or processing may have mediated the transmission of odor preferences to female offspring via yet unknown epigenetic mechanisms. A transmission of acquired pheromone odor preferences may favor assortative mating and chemosensory speciation.

### Conclusion

We have demonstrated the learning, and the inheritance of new behavioral responses to new sex pheromone blends by female *B. anynana* butterflies, calling into question the belief that sexual chemical communication is under stabilizing selection. Over time, as new pheromone blends appear, and new learned sex pheromone preferences for those blends develop, new populations of insects may evolve with specific sensitivity for those blends encoded at the genetic level. Learning to prefer a new sex pheromone blend could be the starting point of the evolution of chemosensory communication, especially if the learned preferences can be inherited.

## Methods

### Husbandry

*Bicyclus anynana* is an African butterfly that produces alternative seasonal phenotypes in response to environmental cues (41). To avoid the seasonal variations in courtship behavior (27), eye size and UV light perception (42), and sex pheromone production (25), we performed all experiments with wet season butterflies, all reared at 27°C, 80% humidity and 12:12h light:dark photoperiod. Larvae were fed young corn plants, and adults mashed banana. Sex was determined at the pupal stage, and females were placed in individual containers stored in a separated incubator, devoid of males or male sex pheromones until a male exposure or a mating trial. Upon emergence, males were put in age-specific cages. They were all naïve, virgin, aged from 4 to 6 days old during the experiment and had dorsal forewing eyespot UV-reflective pupils (as their absence in males is strongly selected against by females (26)). The two males presented to each female for a mating trial had the same age and similar wing size. The experimental procedure is described in Fig. 1.

### Experiment 1: Prevention of MSP2 release from males

Males were prepared following the method described in (28). The ventral hindwing androconia and yellow hair pencil were both coated with transparent non-viscous nail solution (Revlon Liquid Quick Dry). The hindwing dark hair patch, which overlaps the forewing androconia, was left uncoated. This treatment prevents the emission of MSP2 produced by hindwing glands only (24), and causes the reduction of MSP1 and MSP3 total amounts by an average of 70% and 60% respectively (Fig. 1b) (25). The hindwing ventral side of Wild type (Wt1) males received the same treatment to control for the odor of the nail solution. Males were prepared ~16 hours prior to exposure or mate choice trials (Figs. 1a and 1b).

### Experiment 2: Increase of MSP2 amount in males

5µg of MSP2 (Cayman Chemical, n°9001996) diluted in 2µL of hexane were applied to each hindwing androconia of NB2 males. Wild type control males (Wt2) received the same volume of solvent only in the same wing location (Fig. 1c). Hexane was used as a solvent as it didn’t impact naïve female choice (tested in a mate choice assay, described in the Supplementary file 1). The high load of synthetic hexadecanal was chosen to maximize the difference between MSP2 amounts of NB2 and Wt2 butterflies until several hours after application of the solution (Supplementary Fig. 1). The evaporation rate of hexadecanal was determined by gas chromatography from 30 minutes to 8 hours after perfuming. Between perfuming and MSP extraction, two males were placed together in one cylindrical hanging net cage, under identical temperature, humidity and light conditions than the ones used for the mate choice experiment (see “Mate choice assays” below and Supplementary procedure 1). Males were allowed to rest 30 minutes after perfuming until used for exposure or mate choice trials. At the end of this period, NB2 males had similar amounts of MSP2 and MSP3 on their wings (Fig. 1c).

### Female exposure to New Blend or Wild type males

The female butterfly was released in a cylindrical hanging net cage (30cm diameter, 40cm height) less than an hour after emergence (on day 0). The exposure was done manually by retaining the male between the head and the thorax with narrow-tipped featherweight forceps for 3 minutes. The males were presented directly to the females in a similar way as the natural courtship behavior (same distance and orientation). In this position, male fluttering, the first step of the courtship sequence, helped the volatilization of the pheromones and could be encouraged by a gentle squeezing of the forceps (Fig. 1d). This procedure allowed a direct and controlled exposure of the females, and was non-harmful to the males. After exposure, the female remained isolated until day 2, when mate choice assays were conducted (Fig. 1a). Each female (naïve included) was allocated an identification number which doesn’t indicate the treatment she was submitted to, so that the investigator was unaware of the sample group allocation during the mate choice experiment and when assessing its outcome.

### Mate choice assays

All experiments were done at 24°C, 60% humidity, under UV and white light, in cylindrical hanging net cages. Mate choice of naïve and exposed females was started on day 2, around 9:30am (Fig. 1a). One Wt and one NB male were placed in the same cage along with the female. Female’s abdomens were pre-dusted with fluorescent orange powder which is transmitted to the male upon copulation, allowing the identification of the mating partner. Males were checked for presence of powder every 2 hours to prevent multiple mating. Assays were ended after 8 hours after the beginning of the experiment. The latter time point corresponds to MSP2 amounts becoming similar between Wt2 and perfumed males (Supplementary Fig. 1). To differentiate NB2 and Wt2 males, a black dot was applied with a sharpie pen randomly at the top or the bottom of their ventral hindwing. NB1 and Wt1 males were recognizable thanks to the light grey color of the nail solution covering the androconia or the corresponding area of the wing on the opposite side.

### Testing the transgenerational inheritance of mate choice preferences

An additional group of females were exposed to either NB1 or WT1 males, following the same exposure protocol as described above. We didn’t test the preference of offspring of females that choose NB1 males, but instead, each female was mated with a single naïve Wt males in a separate cage. This procedure was followed to prevent possible confounding effects of the mate choice experiment and any predisposed genetic preferences that females may have. The male was removed after mating and the female given a corn plant for egg collection. Each female and its offspring (F1 individuals) constituted a family. F1 pupae were sexed, and the females were submitted to the exact same isolation procedure as naïve females until mate choice assays between a NB1 and a Wt1 male, tested on day 2 using identical procedures as described above (Fig. 1). Around 5 females were tested from the 13 Wt1 and the 11 NB1 families.

### Statistical analyses

Results from experiment 1, including offspring mate choice, and from experiment 2 were analyzed separately using R v. 3.2.4 (43) implemented in RStudio v.1.0.136 (44). P-values were obtained by likelihood ratio tests of full regression models tested against simplified models with specific factors removed.

A Mixed Effect Logistic Regression (MELR) was used to analyze females and their offspring mate choice (experiment 1), as this model includes both fixed and random effects. *Female mate choice* was the binomial response (NB1 male chosen or not). The *family* identity was implemented as a random factor in the model. Each female from the parental generation, taken from our stock population cage, was considered as belonging to different families. The fixed factors included the female *treatment* (NB1-exposed females, offspring of NB1-exposed females, Wt1-exposed females, offspring of Wt1-exposed females, and naïve females) and the *age of males used for mate choice* (4, 5 or 6 days old). The analysis was followed by a pairwise comparison of the significant fixed effects using Tukey Contrasts. Because naïve females (including offspring) were not exposed, the effect of *male age during exposure* (4, 5 or 6 days old) on female choice was analyzed separately with a binomial logistic regression. Packages lme4 (45) and multcomp (46) were used.

The factors that contributed to *female mate choice* in experiment 2 were analyzed with a logistic regression, fitting a Generalized Linear Model (GLM) with quasi-binomial errors to control for over-dispersion, and a logit-link function. Fixed factors used in the model included *treatment* (NB2-exposed females, Wt2-exposed females and naïve females), *male age during exposure* and *male age for mate choice* (in both steps, they were 4, 5 or 6 days old), and the *position of the black mark* used to identify NB2 and Wt2 males (the bottom or the top of the wing).

Finally, in both experiments, actual preference for the NB or the WT blend was tested using a Pearson’s *χ*^2^ test in R. Blends were considered as preferred by females if mate choice differed significantly from random mating [50:50].

## Data availability

The datasets generated during the current study have been submitted to the Institutional repository of the National University of Singapore ScholarBank@NUS (http://scholarbank.nus.edu.sg/).

## Acknowledgment

We thank the National University of Singapore Environmental Research Institute and Frances Lim for the use of the GC-QQQ, Jeremy Pang and Tan Min for performing preliminary experiments, Dr. Erica Westerman, Dr. Marie-Jeanne Holveck, and the butterfly lab members for their help and useful suggestions about the experiments and the manuscript. This work was founded by the Ministry of Education, Singapore grant MOE2014-T2-1-146.

## Author contributions

ED and AM designed the study; ED and LXP performed the experiments and analyzed the data; ED, LXP and AM wrote the manuscript.

## Competing financial interests

The authors declare no competing financial interests.

